# Graph-based algorithms for phase-type distributions

**DOI:** 10.1101/2022.03.12.484077

**Authors:** Tobias Røikjer, Asger Hobolth, Kasper Munch

## Abstract

Phase-type distributions model the time until absorption in continuous or discrete-time Markov chains on a finite state space. The multivariate phase-type distributions have diverse and important applications by modeling rewards accumulated at visited states. However, even moderately sized state spaces make the traditional matrix-based equations computationally infeasible. State spaces of phase-type distributions are often large but sparse, with only a few transitions from a state. This sparseness makes a graph-based representation of the phase-type distribution more natural and efficient than the traditional matrix-based representation. In this paper, we develop graph-based algorithms for analyzing phase-type distributions. In addition to algorithms for state space construction, reward transformation, and moments calculation, we give algorithms for the marginal distribution functions of multivariate phase-type distributions and for the state probability vector of the underlying Markov chains of both time-homogeneous and time-inhomogeneous phase-type distributions. The algorithms are available as a numerically stable and memory-efficient open source software package written in `C` named `ptdalgorithms`. This library exposes all methods in the programming languages `C` and `R`. We compare the running time of `ptdalgorithms` to the fastest tools using a traditional matrix-based formulation. This comparison includes the computation of the probability distribution, which is usually computed by exponentiation of the sub-intensity or sub-transition matrix. We also compare time spent calculating the moments of (multivariate) phase-type distributions usually defined by inversion of the same matrices. The numerical results of our graph-based and traditional matrix-based methods are identical, and our graph-based algorithms are often orders of magnitudes faster. Finally, we demonstrate with a classic problem from population genetics how `ptdalgorithms` serves as a much faster, simpler, and completely general modeling alternative.

## 1 Introduction

A phase-type distribution is the time until absorption in a continuous-time Markov jump process on a finite state space. A discrete phase-type distribution is the time until absorption in a discrete-time Markov chain on a finite state space (Neuts, 1981). Phase-type distributions find many uses in statistical modeling, such as physics, telecommunication, and queuing theory (e.g. Aalen (1995); Faddy and McClean (1999); He (2014); Acal et al. (2019)). Recently, phase-type distributions have been used to model the coalescence process, which describes the genealogical relationship among DNA sequences (Hobolth et al., 2019). The solid theoretical foundation of phase-type distributions (e.g. He (2014) and Bladt and Nielsen (2017)) allows for elegant matrix-based formulations and mathematical proofs. However, computations using matrix-based formulations become infeasible even for systems with just thousands of states. One such example is the basic coalescence model (see Section 3 in Hobolth et al. (2021)), where the number of states grows exponentially fast in the square root of the sample size.

In most models of real-world phenomena, states are sparsely connected, and a graph-based description is thus more natural and efficient than a matrix-based representation. In this article, we use a graph-based approach to develop efficient algorithms for computing the moments, probability distribution, state probability vector, and reward transformations for phase-type distributions. Working with phase-type distributions as graphs has several advantages. First, we can construct the state space iteratively, making it easy to build models for very complex phenomena. Second, we do not store the phase-type distribution as a matrix and thus require much less memory for its representation. Third, the moments and distributions are computed orders of magnitudes faster than matrix operations. These advances are made by constructing algorithms for phase-type distributions that apply directly to the graph structure. We present efficient graph-based algorithms for a generalized iterative state space construction, for reward transformation for positive or zero rewards, for computation of the marginal and joint moments of multivariate phase-type distributions, for computation of the distribution functions, and for computation of the state probability vector of the Markov process at any given time for both time-homogeneous and time-inhomogeneous distributions. The algorithms apply to both continuous and discrete phase-type distributions.

This paper presents theoretical principles and probabilistic interpretation of our algorithms with brief code examples. The algorithms are written in `C` with methods exposed as an open-source library, named `ptdalgorithms`, in `C` and `R`. The library is available on GitHub and is easily installed and compiled using only this `R` code:

~~~
library(devtools)
devtools::install_github("TobiasRoikjer/PtDAlgorithms")
~~~

The full documentation of the package is available at the GitHub website, and as `R` man pages of the `ptdalgorithms R` package.

The paper is organized as follows. We first present the graph-representation of a continuous phase-type distribution (Section 2.1) and show algorithms for constructing the state space (Section 2.2) and for reward transformation in the graph representation (Section 2.3). We then define the moments of phase-type distributions and show how we can compute the expectation, variance, and all higher-order moments recursively on an acyclic graph (Section 2.4). We then show how a normalized form of any cyclic graph can be converted into an equivalent acrylic graph amenable to the recursive computation (Section 2.5 and Section 2.6). We show how the algebraic operations to produce the acyclic form can be cached so that any higher-order moments can be computed in quadratic time (Section 2.7). We then describe our representation of multivariate distributions and how to compute marginal and joint moments of these distributions using rewards (Section 2.8). We go on to show how our framework also accommodates discrete phase-type distributions (Section 2.9), and how their distribution functions can be computed (Section 2.10). We then show how distribution functions for continuous phase-type distributions can be approximated to arbitrary precision using the same approach (Section 2.11). Finally, we show how time-inhomogeneous models can be represented in our framework (Section 2.12). Empirical running times for various uses of `ptdalgorihms` are compared to state-of-the-art matrix based approaches in Section 3. We end the paper by applying `ptdalgorihms` to the isolation-with-migration model from population genetics in Section 4.

## 2 Graph-based algorithms for phase-type distributions

### 2.1 Graph-representation of a phase-type distribution

In our matrix description of a continuous phase-type distribution, we follow Section 3 of Bladt and Nielsen (2017). We assume *p* transient states with the initial distribution *α_i_*, *i* = 1, …, *p,* with a potential defect 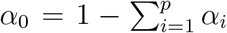. Let ***α*** = (*α*_1_, …, *α_p_*) be the initial distribution vector. The instantaneous sub-intensity matrix (the rate matrix excluding absorbing states) is denoted ***S*** and the Green matrix is ***U*** = (−***S***)^−1^. Entry (*i, j*) in the Green matrix, ***U***_*ij*_, is the expected total time spent in state *j* when the Markov chain begins in state *i*. The cumulative distribution function is given by

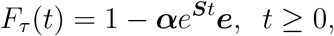

where *e*^***S****t*^ is the matrix exponential of the matrix ***S****t* and ***e*** is the vector of ones.

In our description of phase-type distributions as a weighted directed graph, we follow the standard notation found in e.g. Frydenberg (1990) and Younes and Simmons (2004). A directed graph *G* is a tuple *G* = (*V, E*) where *V* is a set of vertices and *E* is a set of ordered pairs of vertices. An edge from a vertex `v` ∈ *V* to `z` ∈ *V* is denoted (`v` → `z`) ∈ *E*. The set of edges (`v` → `z`) for all `z` ∈ *V* are referred to as the out-going edges of `v` and the set of edges (`u` → `v`) for all `u` ∈ *V* are the in-going edges of `v`. For any pair of vertices connected by an edge (`u` → `v`), `u` is referred to as a *parent* of `v`, and `v` as a *child* of `u`. The set of children of a vertex *v* is denoted `children`(`v`) and the set of parents is denoted `parents`(`v`). Note that if the graph has cycles, a parent of a vertex may also be a child of the same vertex.

A *weighted* directed graph associates each edge with a real-valued weight function *W* : *E* → ℝ that represents the transition rate between two states. The weight of an edge (`v` → `z`) ∈ *E* is denoted *w*(`v` → `z`) ∈ ℝ and the sum of weights of out-going edges for a vertex `v` is denoted

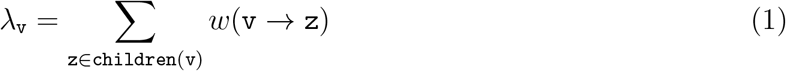

In order for a weighted directed graph to correspond to a valid phase-type distribution defined by the sub-intensity matrix, all weights must be strictly positive real numbers corresponding to exponential rates. Further, edges only connect different vertices (self-loops are not allowed), and at least one vertex must be without out-going edges, representing an absorbing state. Finally, all vertices must have a path to an absorbing vertex with a strictly positive probability.

We designate a single starting vertex denoted as `S` and let its edge weights to other states represent the initial distribution vector such that *w*(`S` → `v`_*i*_) equals *α_i_* and so that *λ*_S_ = 1. A potential defect is modeled by edges going directly to absorbing vertices from the starting vertex. The starting vertex is assigned a reward of zero, as introduced in Section 2.3, so it does not contribute to the time until absorption.

The graph is mutable. We can add vertices, add and remove edges, and change the weight of an edge. We use a prime symbol to denote a transformation or change to a graph, i.e., Applying a set of transformations to *G* gives rise to an updated graph *G*′. This update of *G* is represented as *G* ← *G*′ in the listings of algorithms.

### 2.2 State space construction

We represent the state space as an iteratively constructed graph. Iterative construction greatly simplifies the specification of large and complex state spaces because they can be specified from simple rules governing transitions between states. Our construction algorithm visits each vertex in the graph once and independently of any other vertices. The iterative construction is possible because of the Markov property of phase-type distributions: for any state, all out-going transitions and their rates can be specified knowing only the current state. The construction algorithm is shown in Algorithm 2.1. An example of a state space construction is shown in Figure 1 alongside an example of the `R` code needed to generate the state space.

**Figure 1:**
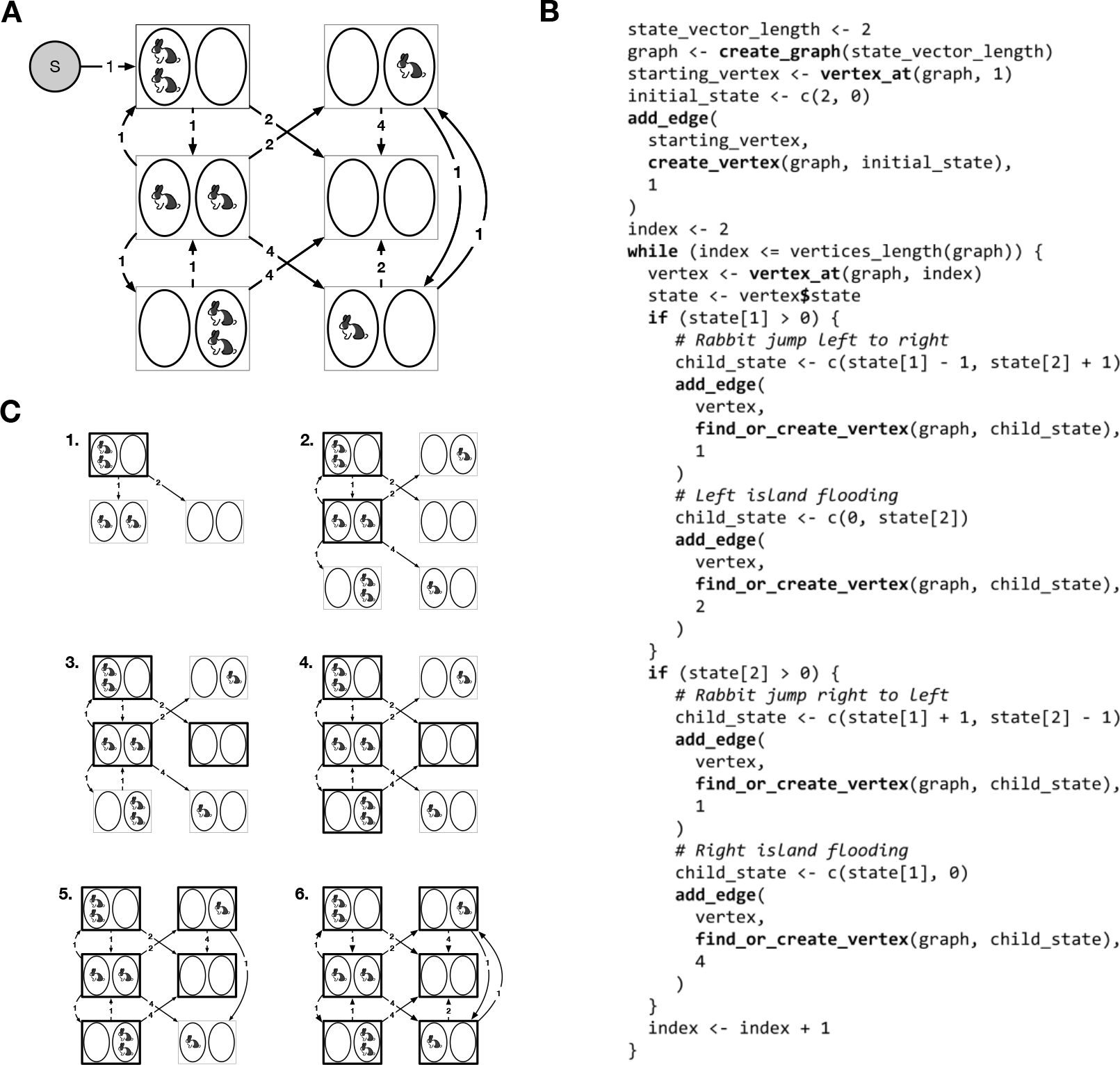
Example of state space construction. In this model, rabbits jump between two islands at a rate of one. The two islands are flooded at rates 2 and 4, drowning all rabbits on the flooded island. The absorbing state is when all rabbits have drowned. **(A)** Shows the state space with transitions. **(B)** Shows the `R` code that generates the state space. States are encoded as vectors of two integers representing the number of rabbits on each island. We construct the state space iteratively by adding vertices and edges. Vertices are visited in the order they are added to the graph. Visited states at each stage are shown with bold outlines. We do not label absorbing states explicitly, as they are simply vertices with no out-going edges. The special starting vertex goes to the initial state with two rabbits on the left island with a probability of one. The graph keeps a record of its vertices and their matching state, and the function `find_or_create_vertex` only creates vertices for states that do not exist. **(C)** The state space after each iteration of the `while` loop in panel B. The `index` variable in panel B enumerates the visited states shown with bold outline, while `vertices_length`(`graph`) returns the number of states currently added to the graph. Once these are equal, the state space construction is completed.

Our algorithm requires an efficient lookup data structure representing a one-to-one bijection between a state and a vertex in the graph. In `ptdalgorithms` we use an AVL tree to map a state to a vertex. We use a vector of integers to describe each state, but any representation of a state with an equivalence relation and an ordering (< relation) can be used.

While the algorithms are independent of the underlying data structure, the asymptotic complexities of our algorithms assume that all children of a vertex `v` can be merged to all of its parents in quadratic time in the number of vertices, either by updating the weight of the edge or by adding a new edge from the parent to the child. This requirement is met if the addition of new edges and updates of weights are both constant-time operations. In `ptdalgorithms`, we store the edges in an ordered linked list, and we can merge two ordered linked lists in linear time by comparing them element-wise in their sorted order. For *O*(|*V*|) parents, we perform *O*(|*V*|) merges each in time *O*(2|*V*|), and therefore in quadratic time, where |*V*| denotes the cardinality (number of entries) of the set *V*.

For modeling convenience, `ptdalgorithms` supports parametrization of edges, allowing edge weights to be updated after the construction of the state space. This feature is useful for exploring the effect of different parameter values in the same model (see `ptdalgorithms` documentation).

To provide an convenient way to inject `ptdalgorithms` into existing workflows, `ptdalgorithms` provides the function `matrix_as_graph`. This function produces a graph from a sub-intensity matrix and an initial probability vector as in this example computing the expectation of the phase-type distribution shown in Figure 1:

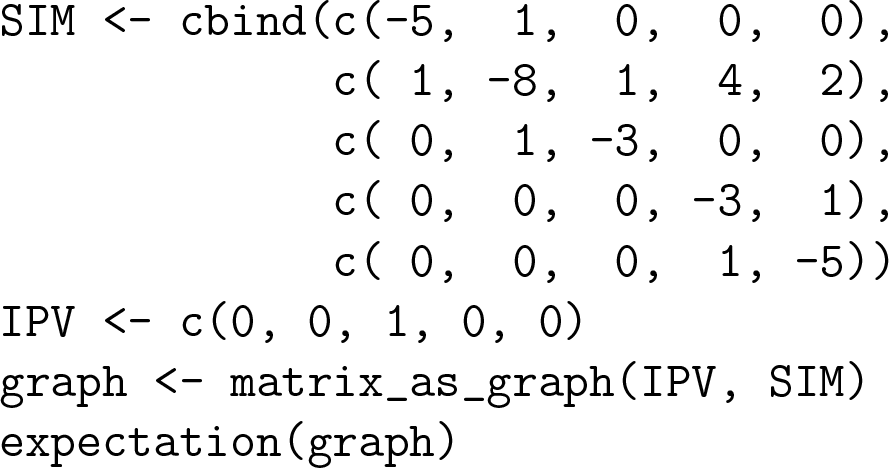

**Algorithm 2.1.**
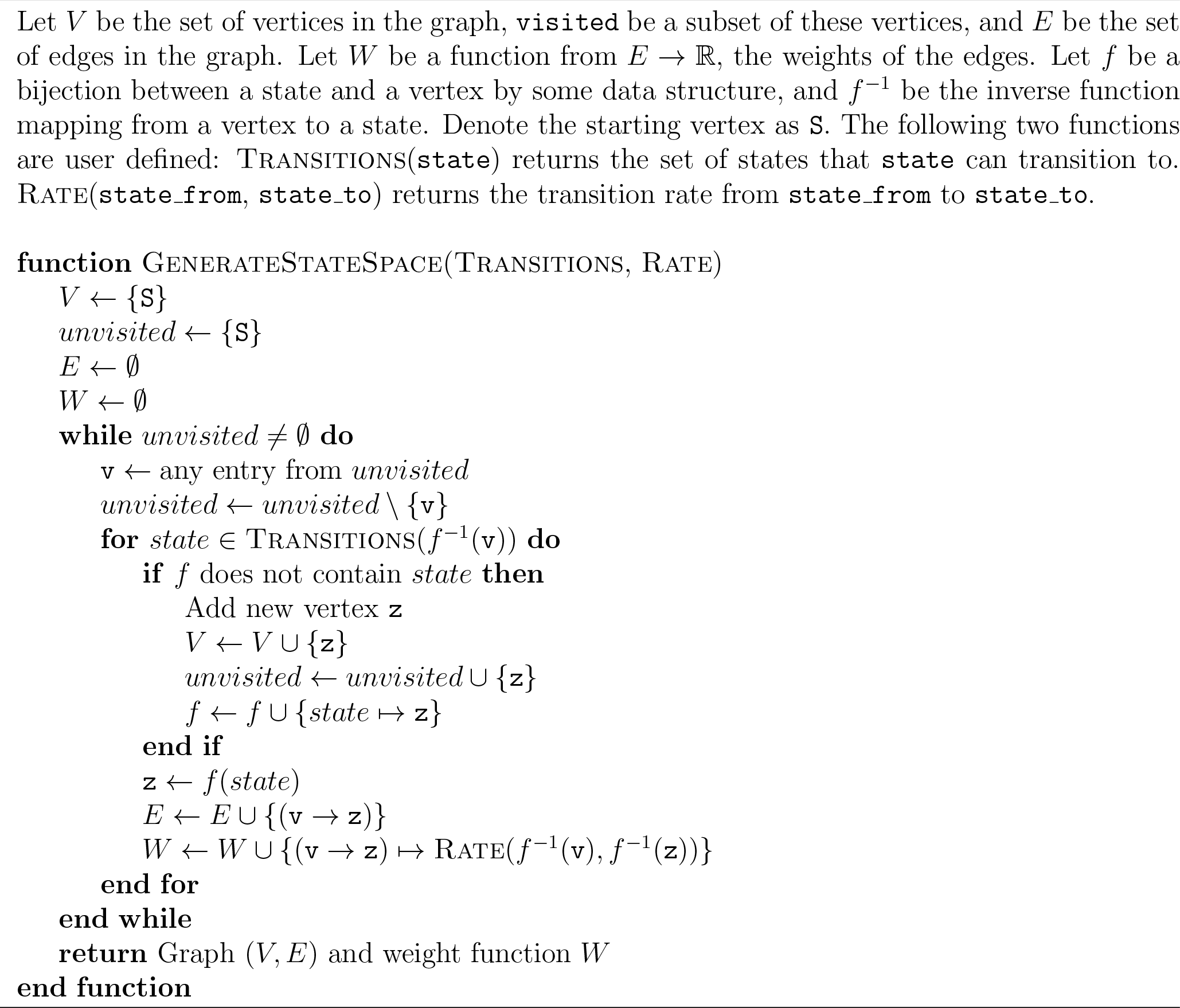
General state space generation algorithm

### 2.3 Reward transformation of a phase-type graph

We can assign a zero or a real-valued strictly positive reward to each state, and the waiting time in each state is then scaled with this reward. The phase-type distribution is no longer the time until absorption but rather the accumulated reward until absorption. If all rewards assigned to each state is one, the phase-type distribution is unchanged. Let *τ* be a phase-type distributed stochastic variable with *p* transient states, *τ* ~ PH_*p*_(***α, S***), and let {*X_t_*} be the underlying Markov jump process. We are interested in the reward-transformed variable

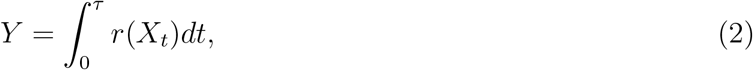

where ***r*** = (*r*(1), …, *r*(*p*)) is the vector of non-negative rewards. Theorem 3.1.33 in Bladt and Nielsen (2017) states that *Y* is also phase-type distributed, and they provide matrix formulas for computing the sub-intensity rate matrix of the reward-transformed variable. In this section, we derive the corresponding graph-based construction.

Consider the reward transformation where a strictly positive reward *r_k_* > 0 is assigned to state *k* and where the rewards assigned to all other states remain unchanged, i.e., *r_i_* = 1, *i* ≠ *k*. The reward-transformed sub-intensity matrix is then obtained by scaling row *k* in the rate matrix by the inverse of *r_k_*

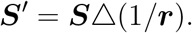

In the directed weighted graph, this corresponds to multiplying the weight of all out-going edges from vertex *k* by 1/*r_k_*.

Now consider the alternative reward transformation with a zero reward *r_k_* = 0 assigned to only state *k* so that *r_i_* = 1, *i* ≠ *k*. In this case, we need to remove the vertex for state *k* from the graph and update the edges and weights accordingly. Before transformation, the transition matrix of the embedded Markov chain embedded has entries

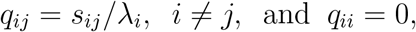

where *s_ij_* is the rate from state *i* to *j*, represented by the weight of edge (*i* → *j*), and where *λ_i_* is the sum of the out-going weights from *i*. The transition probability from state *i* ≠ *k* to state *j* ≠ {*k, i*} for the embedded Markov chain with state *k* removed is

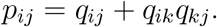

The waiting time in each state of the reward-transformed Markov jump process must remain the same, and therefore the intensity from state *i* to state *j* must be 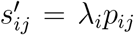. The sub-intensity matrix for the Markov jump process with state *k* removed therefore has entries

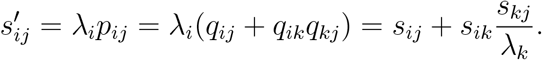

In terms of graph-operations this means that if edge (*i* → *j*) already exists (if *s_ij_* > 0), we should add *s_ik_s_kj_*/λ_*k*_ to the weight of that edge. If the graph does not have edge (*i* → *j*) already (if *s_ij_* = 0), then we add it to the graph with weight *s_ik_s_kj_*/λ_*k*_. The resulting algorithm is shown in Algorithm 2.2 and an example of a reward transformation is shown in Figure 2.

**Figure 2:**
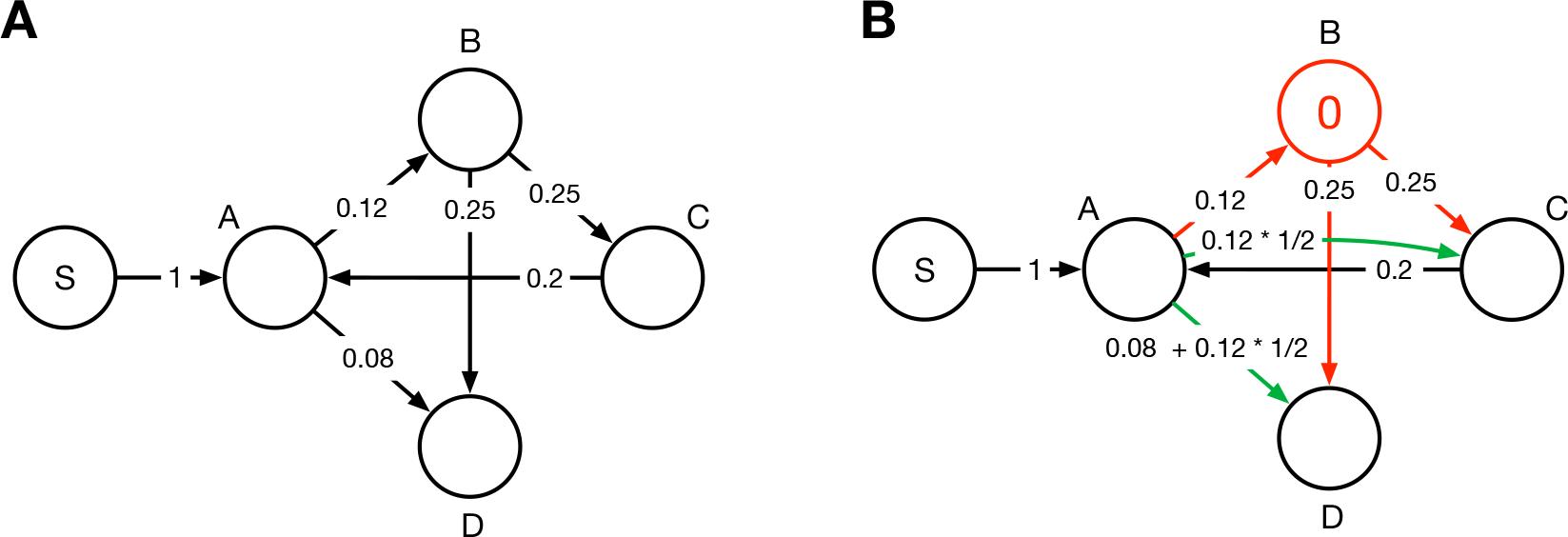
Reward transformation with zero reward. **(A)** Original graph. **(B)** Graph after reward transformation assigning a zero reward to state `B` and a reward of one to states `A`, `C` and `D` in accordance with Algorithm 2.2. Red vertices and edges are removed in the transformation. Green edges are added or updated.

**Algorithm 2.2.**
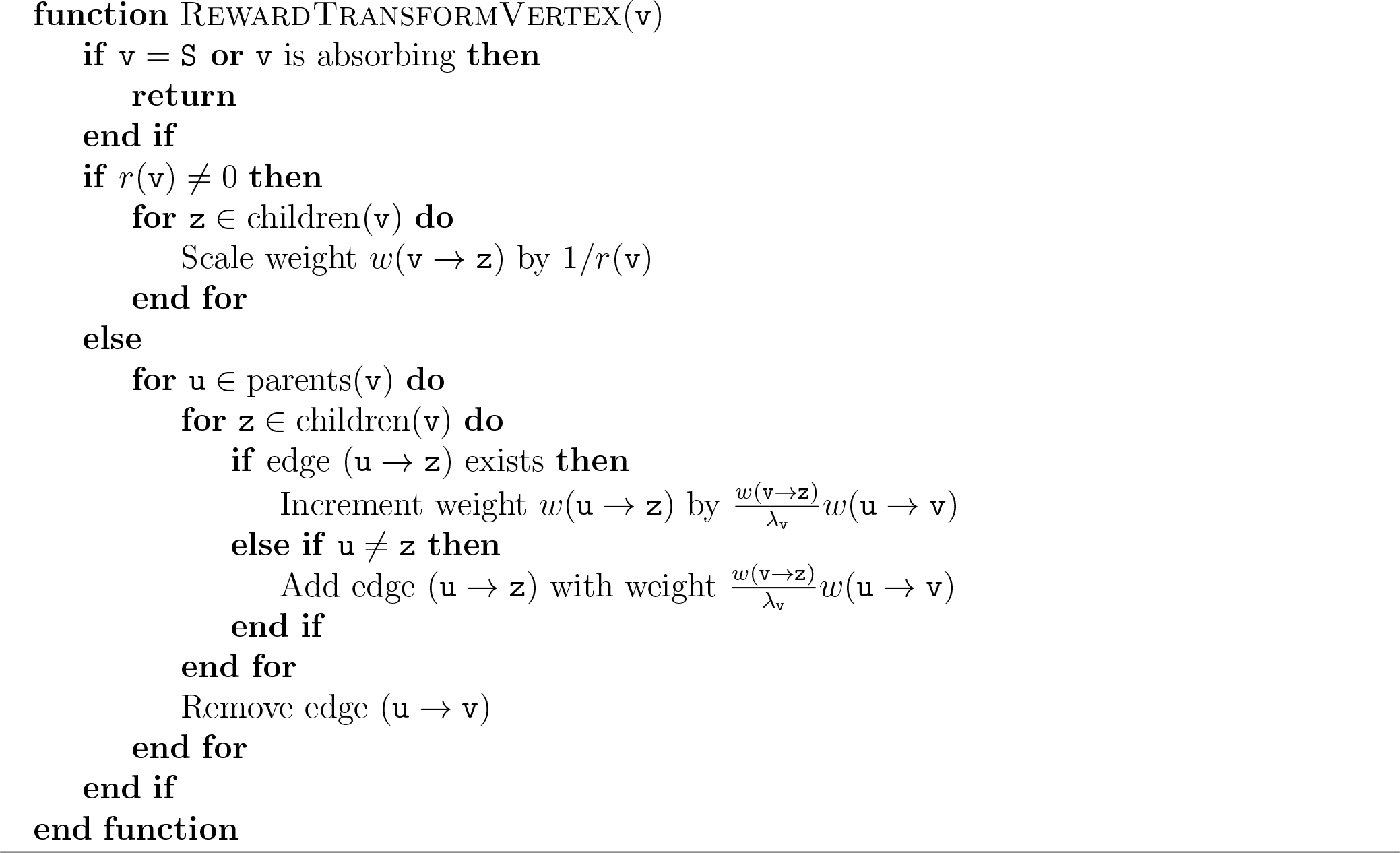
Reward transformation in a graph

### 2.4 Properties of higher-order moments

The higher-order moments of reward-transformed phase-type distributions are determined by matrix-based equations (e.g., Theorem 8.1.5, Bladt and Nielsen (2017))

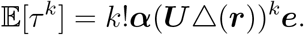

The expectation is thus

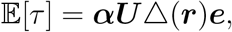

and the second moment is

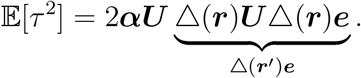

By the associative property of matrix multiplication, we can first compute the part △(***r***)***U***△(***r***)***e***, which gives us a column vector that we can write as △(***r′***)***e***. This property implies that using a different reward vector, the second moment can be computed the same way as the first moment. By induction, we find a new set of rewards such that:

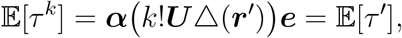

where *τ*′ is a new phase-type distribution with the same state space as *τ*, but with different rewards. In the sections below, we will use this property to construct an algorithm for all moments as well as an algorithm for all joint moments.

### 2.5 A graph representation of normalized phase-type distributions

We can normalize any rewarded phase-type distribution so that intensities for each state sum to one. This is done by adjusting each reward so that the reward divided by the total intensity remains the same after normalization. In doing so, state intensities become transition probabilities that expose the embedded Markov process. In this form, the expected accumulated reward at each state is simply the expected number of visits to each state scaled by its reward. In the sections below, we will represent phase-type distributions in this way to provide simpler proofs and algorithms with probabilistic interpretations.

In the standard graph-based formulation of rewarded phase-type distributions, both the reward and the transition rate contribute to the weight of each edge. In our representation of the *normalized* distribution, we factorize each edge weight into the exponentially distributed expected waiting time at each parent vertex and the remaining transition probability. In the graph, we attribute each vertex `v` ∈ *V* with a scalar 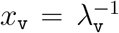 that represents the expected waiting time. This leaves the weights of out-going edges to represent transition probabilities, which sum to one. In the resulting *normalized* graph, the total intensity at each vertex `v` ∈ *V* that is not absorbing is unchanged and expressed as

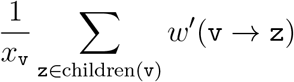

where *w*′(`v` → `z`) = *w*(`v` → `z`)*/λ*_v_. At the starting vertex, *x* is set to one, and for the absorbing vertices, it is set to zero.

Acyclic phase-type distributions often appear and constitute an important special case where the sub-intensity matrix can be reorganized into an upper triangular matrix (Cumani, 1982). In our graph formulation, such graphs have a topological ordering, which allows us to compute the expectations by simple recursions. Let `v` and `z` be vertices and let 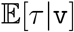 be the expected accumulated reward until absorption given a start at the vertex `v`. For the normalized graph, this gives us the recursion known from the first-step analysis of Markov chains:

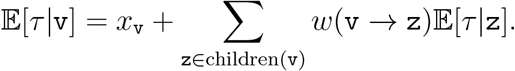

Since we have a single starting vertex, the expectation of the phase-type distribution is

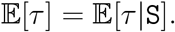

The topological ordering of vertices allows the recursion to be computed using dynamic programming, and we can compute the *k*’th moment in quadratic time in the number of vertices *O*(|*V*|^2^*k*). This is not possible if the state space has cycles since the recursion will never finish. However, as we show below, we can transform the graph for any cyclic phase-type distribution into a graph for an acyclic phase-type distribution with the same states.

### 2.6 An acyclic graph representation of phase-type distributions

We can manipulate the edges *V* and vertex scalars *x* in the graph for a cyclic phase-type distribution to produce an acyclic graph, and we can do this in such a way that the expected time (or accumulated reward) until absorption remains the same. Once constructed, this acyclic graph allows the expectation to be computed recursively as described in the previous section. Algebraically, we find a phase-type distribution such that

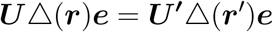

where ***S***′ = (−***U***′)^−1^ constitutes an upper-triangular matrix. This matrix can be found by Gaussian elimination of the system of linear equations expressed as −***Se*** = △(***r***)***e***. Gaussian elimination of sparse matrices by graph theory is a well-studied problem (e.g. Duff et al. (2017)). In `ptdalgorithms` we apply this technique directly on the graph. We will assume below that the phase-type distribution has been normalized as described in Section 2.5.

We first index each vertex arbitrarily to {1, 2, …, |*V*|}, with the only constraints that the starting vertex `S` has index 1 and the absorbing vertices have the largest indices. We then visit all vertices in indexing order. The initial graph is denoted *G*^(0)^ and after visiting the vertex with index *i*, the graph is referred to as *G*^(*i*)^. The algorithm ensures three invariants after each vertex *i* is visited:

1. The vertex has no in-going edges from vertices with a higher index.
2. The expected accumulated rewards until absorption, starting at any vertex, is preserved.
3. The sum of weights for all out-going edges remains 1 (but 0 for the absorbing vertex). In the matrix formulation, this means that

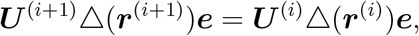

where ***U***^(*i*)^ is the Green matrix for the phase-type distribution given by the graph *G*^(*i*)^, and ***r***^(*i*)^ are new associated rewards. As vertex *i* has no in-going edges from any vertex *j* where *j* > *i*, the graph has a topological ordering and is acyclic once all vertices have been visited. We show in a later section that we only need to perform this relatively expensive computation once in order to compute any moment or joint moment.

We first show how we can remove an in-going and an out-going edge of a vertex without changing the expectation. Let `v` and `z` describe vertices with an associated index. Let 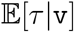 be the expected accumulated reward until absorption given that we start at vertex `v`. The recursion to compute this expectation, as described in Section 2.4, applies but will not end for a graph with cycles

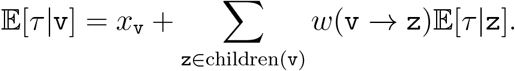

However, we can expand the recursion by bridging the immediate child:

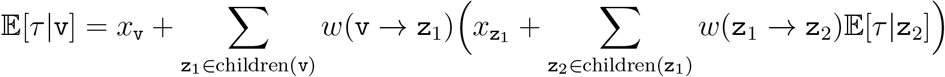

and produce the equivalent equation

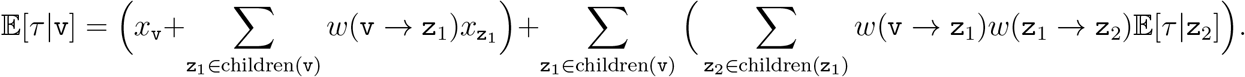

This reveals that we can remove the edge (`v` → `z`_1_) without changing 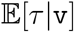 if we apply two operations: we increase the expected waiting time *x*_v_ to 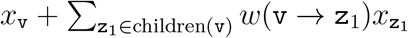 and we change weights of all edges *w*(`v` → `z`_2_) for all `z`_2_ ∈ children(`z`_1_) to *w*(`v` → `z`_1_)*w*(`z`_1_ → `z`_2_).

I.e., if we want to remove `z`, we increase the expected waiting time of any parents by the ingoing weight multiplied by *x*_z_, and add or update edges from each parent to all children of `z` by the product of the edge weights. These two operations allow us to remove vertex `z` from a state-path without changing the expectation.

In the situation where vertex `v` is also a child of vertex `z`, the above procedure creates a self-loop that needs to be resolved: a self-loop with weight *w*(`v` → `v`) can be removed without changing the expectation if we increase *x*_v_ to 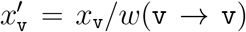, and scale all other out-going edges by 1/(1 − *w*(`v` → `v`)).

By our three invariants, once we visit the vertex with index *i*, `v`_*i*_, all children of vertex `v`_*i*_ will have a higher index because all vertices with indexes lower than *i* have been visited. At this point, not all parents of `v`_*i*_ have an index smaller than *i*. However, redirecting edges from parents with a larger index establishes the three invariants for vertex *i*. The equation for 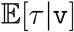 above shows that the expected accumulated reward until absorption is unchanged, and by induction, an acyclic graph is produced once all vertices are visited. The procedure is summarized in Algorithm 2.3 and lets us build an acyclic graph in *O*(|*V*|^3^) time. A worked example of the algorithm is shown in Figure 3.

**Figure 3:**
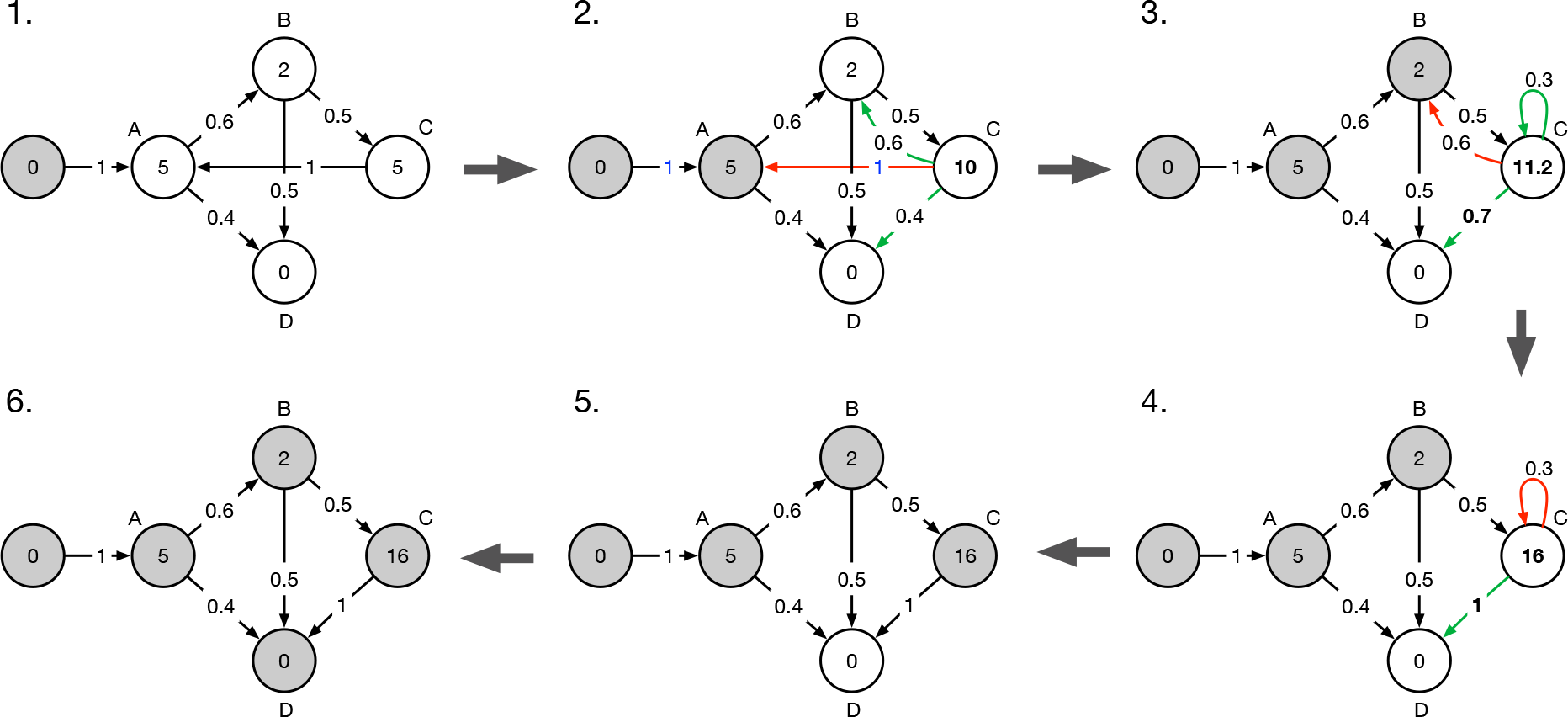
Ayclic graph construction. Example conversion of a normalized graph to acyclic form using Algorithm 2.3. Visited vertices are colored grey. Removed edges are colored red and new or updated edges are colored green. The saved parameterized vertex updates are as follows. Step 2: *x_C_* ← *x_C_* + *x_A_*. Step 3: *x_C_* ← *x_C_* + *x_B_* · 0.6. Step 4: *x_C_* ← *x_C_* + *x_C_* · (1/(1 − 0.3) − 1).

The algorithm can now compute the expected accumulated reward recursively using dynamic programming in *O*(|*V*|^2^) time on the acyclic graph. We show in section section 2.7 that we only need to construct the new acyclic graph once. In `ptdalgorithms` we improve the empirical running time of the acyclic construction by indexing vertices by topological ordering if one exists or by some ordering by the strongly connected components.

To see the correspondence between the acyclic graph construction and Gauss elimination, note how the system of equations −***Se*** = △(***r***)***e*** corresponds to the example graph show in Figure 3 when

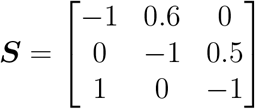

and ***r*** = (5, 2, 5). We can write the system of equations as:

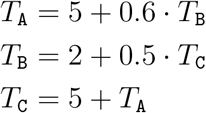

where 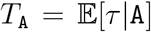 is the expectation starting in state `A` and similarly for *T*_B_ and *T*_C_. The zero expectation starting at the absorbing state, *T*_D_, does not appear in the equations. Gauss elimination makes use of three different operations. The first is ”Swap positions of rows”. In our algorithm the appropriate ordering is maintained by visiting vertices in index order. The second is ”Add to one row a scalar multiple of another” (insert one equation in another). This corresponds to the graph operations removing an edge to a parent with higher index. For example, in Step 1 in Figure 3, the removal of edge (`C` → `A`) and updates of edges (`C` → `B`) and (`C` → `D`) corresponds to inserting the first equation in the third. The third operation is ”Multiply a row by a non-zero scalar” (remove multiple instances of a variable in an equation). This corresponds to the graph operations removing a self-loop. For example, in Step 4 in Figure 3, the removal of self-loop (`C` → `C`) and update of edge (`C` → `D`) corresponds to isolating *T_C_* in the equation *T*_C_ = 11.2 + 0.3 · *T*_C_. Once all vertices have been visited and the graph is acyclic, the system has an upper triangular form

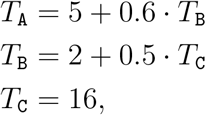

and can be solved by back-substitution. On the graph this corresponds to the recursion in topological order at the end of Algorithm 2.3.

### 2.7 Computing higher-order moments in quadratic time

In the conversion to an acyclic graph, the scalars *x* associated with vertices are updated in a series of increments. In the implementation of Algorithm 2.3, we save this list of updates rather than applying them directly. The resulting list of update functions is at most *O*(|*V*|^2^) long as we visit each vertex at most once and update the scalars of at most |*V*| parents. We also save the update functions used to compute the expectation on the acyclic graph using dynamic programming (also *O*(|*V*|^2^) long). The two list of instructions compute the expectation in *O*(|*V*|^2^) time.

The complexity of identifying the operations required to convert a cyclic graph to an acyclic graph is *O*(|*V*|^3^). However, only *O*(|*V*|^2^) updates of the scalars *x* are required and the edge weights of the resulting acyclic graph are independent of the expected waiting times at each vertex. This means that once the normalized acyclic graph is constructed and its edge weights are known, it can be re-constructed for an alternative set of rewards using only *O*(|*V*|^2^) updates of scalars *x*. Because higher-order moments of phase-type distributions are just expectations with different rewards, as described in Section 2.4, we can compute any number of moments in *O*(|*V*|^2^) time once the acyclic graph has been constructed. In the `ptdalgorithms` library, the acyclic graph and list of update functions are only created the first time the user calls a moment function, after which all subsequent moment computations run in *O*(|*V*|^2^) time.

### 2.8 Multivariate distributions

Instead of assigning a single real-valued reward to each state, we can assign a vector of real-valued rewards. The outcome of the phase-type distribution is now a vector of positive real numbers. This multivariate phase-type distribution is defined in Chapter 8 of Bladt and Nielsen (2017) as

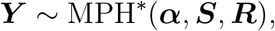

where ***R*** is now a matrix of rewards, such that each row is the accumulated reward earned at the state with that index. A single column of ***R*** represents a univariate phase-type distribution as described above. This means that the marginal moments of a multivariate phase-type distribution can be computed using the graph algorithms already described and still require only a single computation of the acyclic graph.

The joint distribution of a multivariate phase-type distribution represents the conditional outcome of marginal phase-type distributions. Joint moments are also well defined as matrix-based formulations (Theorem 8.1.5 Bladt and Nielsen (2017)). The first cross-moment is

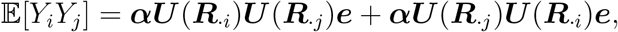

and thereby the covariance is

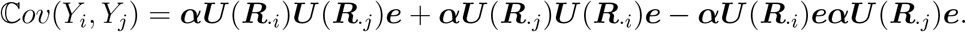

As for all other moments, the joint moments can be computed in quadratic time after construction of the acyclic grap. Computing the covariance matrix of *ℓ* rewards can thus be done in *O*(|*V*|^3^ + *ℓ*^2^|*V*|^2^) time. The utility functions `expectation, variance, covariance`, and `moments` functions provided by `ptdalgorithms` support multivariate phase-type distributions.

**Algorithm 2.3.**
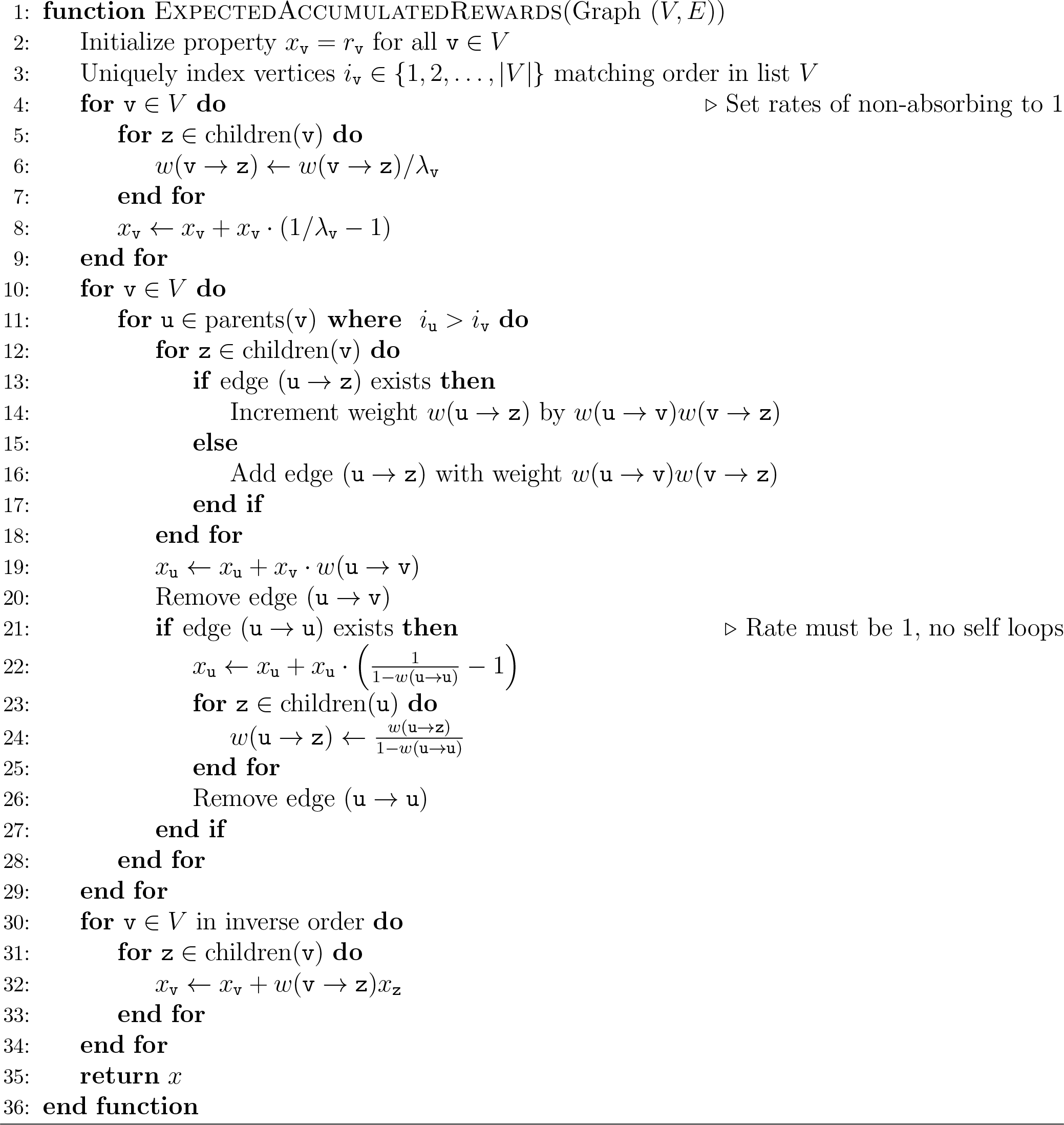
Expected accumulated rewards until absorption

### 2.9 Discrete phase-type distributions

A discrete phase-type distribution is the number of jumps in a discrete Markov chain until the absorbing state is entered. States have a total transition probability of one, and the state space is traditionally represented by a sub-transition matrix ***T*** of rates between non-absorbing states. The initial probability vector is ***π***. As for the continuous phase-type distributions, the expectation is 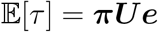, and here the Green matrix is given by ***U*** = −(***T*** − ***I***)^−1^.

Our graph representation for continuous phase-type distributions directly accommodates unrewarded discrete phase-type distributions. We do not represent self-transitions as self-loops, (`v` → `v`), as these are not compatible with our graph algorithms. Instead, we represent self-transitions by the missing transition rate 1 − *λ*_v_. After normalization of the graph, the sum of out-going weights is one, and each vertex scalar *x*_v_ is equal to the geometric expectation of consecutive visits to the state (i.e., the transition to v and immediate self-transitions).

In the normalized discrete phase-type distribution, the sub-transition matrix ***T*** has a diagonal of zero, and ***T − I*** thus shares the properties of the normalized sub-intensity matrix ***S***: the diagonal entries are −1 and the row sums for non-absorbing states is zero. We can thus apply our moment Algorithm 2.3 to discrete phase-type distributions.

Rewarded discrete phase-type distributions and multivariate discrete phase-type distributions have been described thoroughly in Navarro (2019). We can translate the matrix-based reward transformation algorithm from Theorem 5.2 in Navarro (2019) into one operating on the graph representation. Consider a vertex `v` with reward *r*_v_ ∈ ℕ. To represent this integer reward, we augment the graph with a new sequence of connected auxiliary vertices, 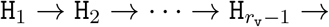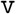, each connected by a single edge with weight 1. The last auxiliary vertex has an edge with weight 1 to the vertex `v` and the edge (`v` → `H`_1_) has weight equal to the self-transition rate. By further redirecting all in-going edges from `v` to `H`_1_ instead, we ensure that each visit to the vertex `v` results in *r*_v_ jumps in the unrewarded discrete phase-type distribution. Reward transformation of zero rewards is achieved using the algorithm for the continuous phase-type distribution.

Higher-order moments are well defined for rewarded discrete phase-type distributions (see Proposition 5.7 in Navarro (2019). The first moment is

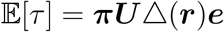

and the second moment is

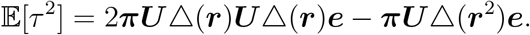

We can construct a multivariate discrete phase-type distribution by associating a vector of zero or positive integers as rewards to each vertex. The moment generating function is well defined for multivariate discrete phase-type distributions by matrix-equations (Section 5.2.4 in Navarro (2019)). For example, the first cross moment is

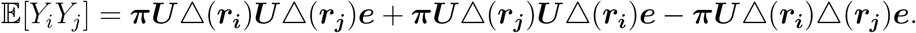

Although the moments of discrete and continuous phase-type distributions are defined differently, terms involving ***U*** still have the form ***U***△(***r***)***e*** from the right-hand side, which reduces to a single vector of rewards ***U***△(***r***)***e*** = △(***r′***)***e*** = ***r′***. These rewards correspond to the row sums of the Green matrix computed for the previous moment in the same way as described for continuous phase-type distributions in Section 2.4. This allows the *O*(|*V*|^2^) computation of higher order moments as described for continuous phase-type distributions in Section 2.7.

### 2.10 Distribution function of discrete phase-type distributions

The probability mass function (PMF) of a discrete phase-type distribution at time *t* describes the probability of the Markov chain entering the absorbing state exactly at jump *t*. A defective distribution, i.e. one where the initial distribution vector does not sum to 1, will have a non-zero probability at *t* = 0. Likewise, the cumulative distribution function (CDF) describes, for time *t*, the probability that the Markov chain has entered the absorbing vertex at jump *t* or before. In matrix form, the CDF is given by

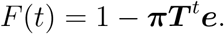

We can express the PMF of the discrete phase-type distribution as a recursion in *t* (Eisele, 2006). Using the law of total probability we get

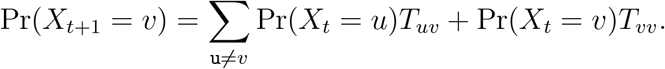

In our graph, we represent the probability of staying at vertex `v` after *t* jumps as a vertex-property `v`*.p*[*t*] and compute the recursion as the probability that a parent will transition to the vertex in the next jump, or that the vertex will jump to itself by a self-loop

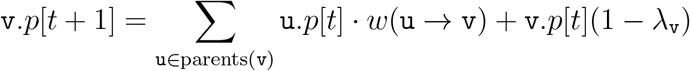

for *t* ≥ 0.

We define the base-case at *t* = −1. In the base-case, `v`*.p* is zero for all vertices except the starting vertex S where it is one. Using dynamic programming, we can find the PMF (and thereby the CDF) at time *t* by first computing the PMF at time 0, 1, …, *t* − 1. To compute the CDF at time *t*, we sum over the PMF at times 0, 1*, …, t*. The asymptotic complexity of the computation to time *t* is *O*(*t*|*V*|^2^). For sparse matrices, or for a relatively low number of jumps (*t* ≪ |*V*|), this algorithm is empirically very fast as shown in Figure 4 and serves as an efficient alternative to the matrix formulation. The algorithm for computing the CDF is shown in Algorithm 2.4.

**Figure 4:**
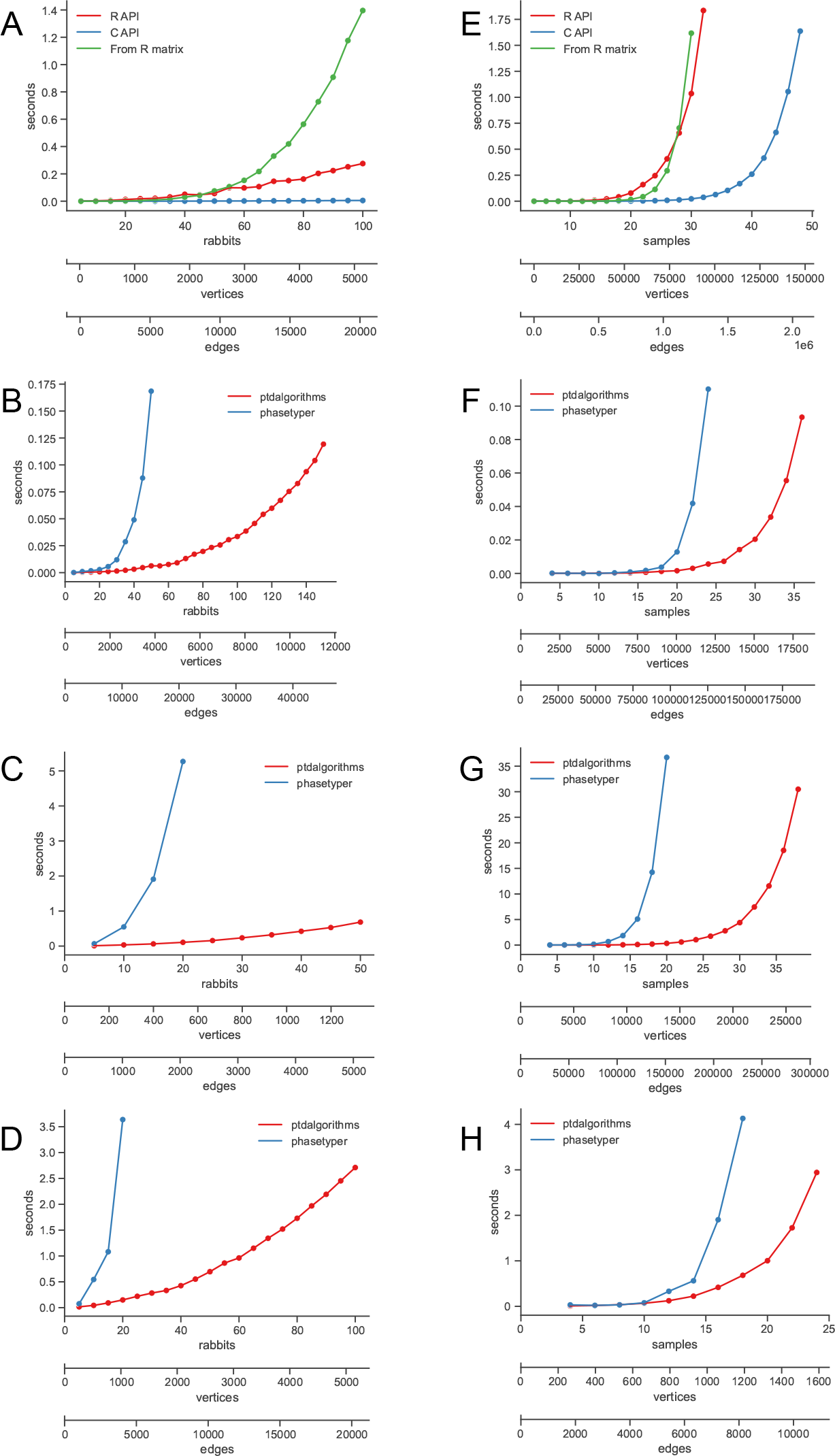
Empirical running time experiments. **(A)** and **(E)**: Time (in seconds) it takes to construct the rabbit and coalescent models for different numbers of rabbits and DNA samples. Each panel also shows the number of vertices and edges in the constructed graph. **(B)** and **(F)**: Time it takes to compute the expectation. **(C)** and **(G)**: Time to compute 100 higher moments or cross moments. **(D)** and **(H)**: Time it takes to compute the CDF for 100 evenly spaced values up to a cumulative probability of 0.95 using a granularity of 10000.

### 2.11 Distribution function for continuous phase-type distributions

In the matrix formulation of continuous phase-type distributions, computing the CDF requires exponentiation of the sub-intensity matrix. However, a continuous phase-type distribution can be approximated, with arbitrary precision, by a discrete phase-type distribution (Bobbio et al., 2004). The number of discrete steps occupying each state has a geometric expectation approximating the exponential expectation of the continuous distribution. This allows us to efficiently compute the PDF, the CDF, and the probability of occupying each state at any time using Algorithm 2.4.

The precision is determined by the number of discrete steps per unit of time, which in turn is controlled by a granularity parameter in `ptdalgorithms`. A granularity of 1000 implies that each time unit of the continuous distribution is divided into 1000 discrete steps. As the graph representations we use for discrete and continuous phase-type distributions are identical, and because self-loops are represented by the remaining transition probability, we can simply divide all out-going weights by this granularity (except for the weights of the starting vertex). The only constraint is that the granularity must be at least as large as the largest transition rate so that the out-going rate of all states are smaller than one after dividing by the granularity. In `ptdalgorithms` the granularity can be set by the user to control the precision of the approximation, but defaults to a value that is at least two times the highest out-going rate of a vertex, and always at least 1024.

**Algorithm 2.4.**
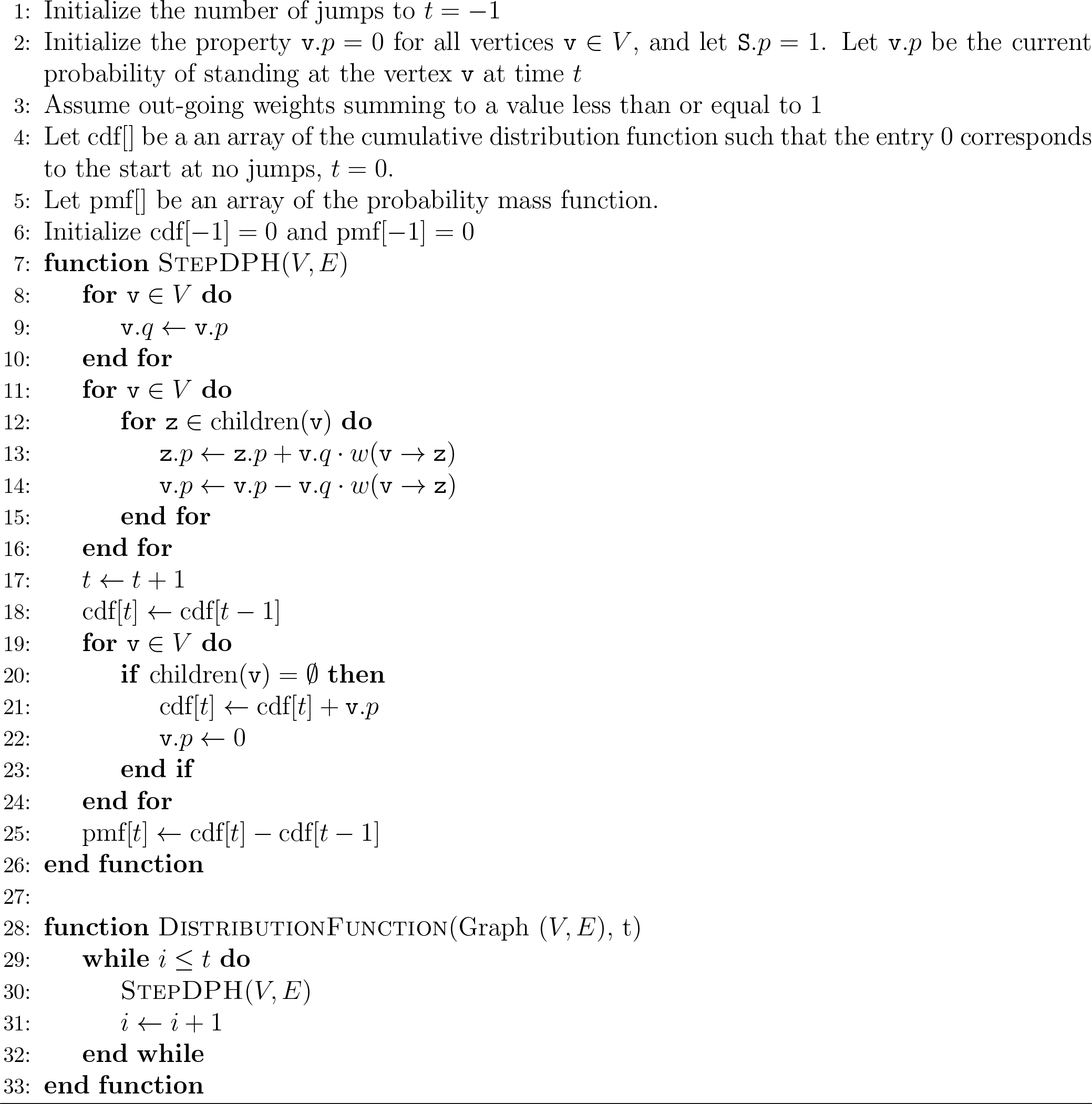
The distribution function of a discrete phase-type distribution

To verify the numerical accuracy of this approach, we compared to the matrix exponential of a phase-type distribution with 1000 fully connected states. Transition rates were sampled randomly in the interval [0, 1], and each graph vertex thus had an average total out-going rate of 500. With a granularity of 10,000, the average self-loop probability is thus 0.95. We computed the cumulative distribution function for times (0, 0.01, 0.02, …, 1.00) using both Algorithm 2.4 as implemented in the `ptdalgorithms` package and using the matrix exponential 1 − ***α****e*^***S****t*^***e***. The average absolute and maximum differences were 0.00003 and 0.0002, which demonstrates stability and a very small negligible numerical difference between the two approaches.

### 2.12 Time-inhomogeneous discrete phase-type distributions

So far, the algorithms have assumed that the phase-type distributions are time-homogeneous, i.e., that the rates between states are constant in time. The alternative time-inhomogeneous phase-type distributions are also well-described using matrix-based equations (Albrecher and Bladt, 2019). Although time-inhomogeneous phase-type distributions are not the focus of this paper, we note that our algorithm for computing the distribution function (Algorithm 2.4) can be applied to produce the distribution and state probability vector of a time-inhomogeneous discrete and continuous phase-type distributions as well. This is achieved by changing edge weights at each time step (e.g. using the function `graph_update_weights_parameterized` in `ptdalgorithms`) in effect allowing the edge weight or the existence of an edge to be a time-dependent variable. Having computed the probability distribution in this way, we can compute all moments of unrewarded time-inhomogeneous phase-type distributions by integration. We can also compute the expectation of a rewarded time-inhomogeneous phase-type distribution by summing the expected visiting time multiplied by the reward for each vertex.

Because Algorithm 2.4 computes the probability of standing at each vertex at any given time step, we can compute the accumulated waiting time in each state ***a***(*t*) at some time *t*. This gives us an alternative way to compute the expectation of a reward-transformed truncated distribution as the dot product ***r · a***(*t*). In models where the state space changes only at one or a few points in time, this provides an efficient means to compute the expectation as the sum of expectations of truncated time-homogeneous distributions.

## 3 Empirical running time

Here we describe the empirical running time of state space construction, moment computation, and computation of the cumulative distribution in the rabbit island model shown in Figure 1 as well as in the coalescent, a standard model in population genetics. The coalescent describes the decreasing number of ancestors to a sample of DNA sequences as time progresses into the past (Kingman, 1982). All experiments are done on a MacBook Pro with an i7 processor using a single core.

Although all features of the `ptdalgorithms` library are exposed as `R` functions, the similar `C` API offers an efficient alternative for the generation of very large state spaces. In Figure 4A and Figure 4E, we show the time it takes to construct the two state spaces for different numbers of rabbits and DNA samples using both the `R` and `C` APIs. For example, for 1000 rabbits, where the system has more than half a million states, we can construct the state space and compute the expectation in two minutes (not shown).

We compare the running time of `ptdalgorithms` to that of `PhaseTypeR` (Rivas-González et al., 2022), which implements the same functionality as `ptdalgorithms` using matrix computations. `PhaseTypeR` uses `expm::expm()` (Goulet et al., 2021) for exponentiation. We have evaluated the algorithms for exponentiation implemented in `expm` and use `PadeRBS` which performs well for both models. We further modified `PhaseTypeR` so that it uses the `matrixdist R` package, which wraps the C++ Amadillo library, for matrix inversion. To our knowledge, this makes `PhaseTypeR` the fastest available matrix-based approach to phase-type distributions. The plots in Figure 4 show the time `ptdalgorithms` and `PhaseTypeR` use to compute the expectation, 100 higher moments or cross-moments (e.g., a 10×10 covariance matrix) and the CDF for 100 evenly spaced values up to a cumulative probability of 0.95. Panels B-D and F-H show the computation times for the rabbit model and the coalescent model.

## 4 An application in population genetics

To demonstrate the capabilities of `ptdalgorithms`, we feature an example from the field of population genetics. Kern and Hey (2017) describe and implement a method for the computing the joint site frequency spectrum (JSFS) for an isolation-with-migration (IM) model. A site frequency spectrum (SFS) is the number of single-base genetic variants shared by 1, 2, …, *n* − 1 sequences among the *n* sequences sampled from a population. The JSFS tabulates the number of variants shared by *i* sequences in one population and *j* in another. The JSFS matrix can be obtained empirically and is determined by the population sizes of the two populations and the common ancestral population, the migration rate between the two populations, and the time that the two populations have been separated (see Figure 5A).

**Figure 5:**
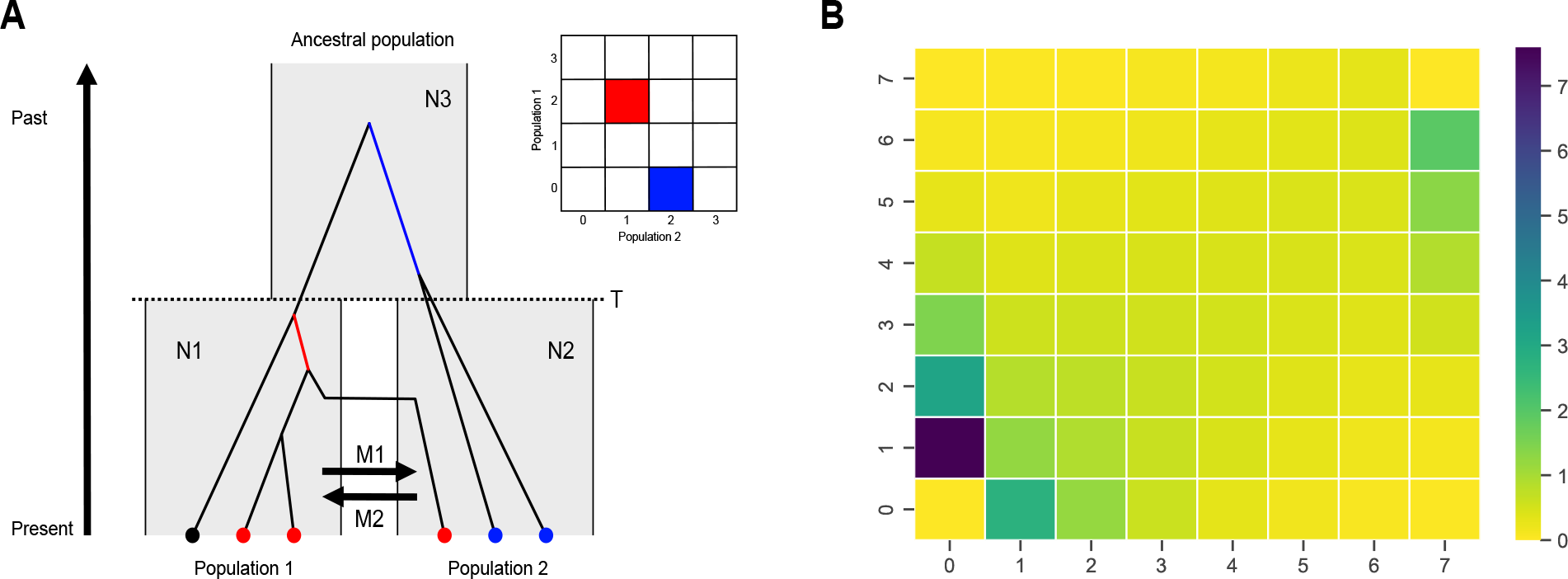
The exact joint site frequency spectrum of an isolation-with-migration model. **(A)**: The isolation-with-migration model. One ancestral population is split in two at time *T*. Before time *T* the two descendant populations are only connected by migration at rates *M*1 and *M*2 in each direction. Scaled population sizes are *N*1, *N*2, and *N*3. In the example genealogy, the red branch has two descendants in population one and one descendant in population two. The blue branch has two descendants in population two and none in population one. The insert matrix shows the entries in the JSFS that the two example branches contribute to. **(B)**: Exact joint site frequency spectrum. Population-scaled parameters are N1=2, N2=1, N3=4, M1=0.005, M2=2, T=3). Each (*i, j*) cell shows the expected length of branches with *i* descendants from population one and *j* descendants from population two.

A genetic variant that appears in three samples from population one and four samples from population two arose from a mutation on a genealogical lineage with three descendants in population one and four descendants in population two. Knowing the mutation rate, the JSFS is given by the expected length of branches with different numbers of descendants in each population.

The structured coalescent models the length of the ancestral branches, and formulated as a continuous phase-type distribution, states may represent ancestral lineages with particular numbers (*i, j*) of descendants in each population. The initial state represents *i* and *j* present-day individual lineages, each with a single descendant. The absorbing state is the state where all lineages have found a single common ancestor. The time of absorption corresponds to time to the most recent common ancestor (TMRCA).

The IM model is not time-homogeneous as it changes from two separate populations to a shared ancestral population at a time *T*. Using `ptdalgorithms`, we can easily model this with one time-homogeneous continuous phase-type distribution truncated at time *T* and another time-homogeneous distribution representing the system after time *T*, as described in section 2.12.

The expected length of genealogical branches with *i* and *j* descendants from the two populations is readily computed after appropriately reward transforming the two phase-type distributions. With seven samples from each population, the two state spaces have 123,135 and 2,999 states.

Kern and Hey (2017) computed the JSFS in around 15 minutes using 12 cores. For comparison, the state space construction and computation of JSFS takes only 35 seconds on a single core using `ptdalgorithms` (Figure 5B). This is particularly noteworthy because `ptdalgorithms` is a general purpose library not tailored to this specific problem. The code for construction of the model and computing the JSFS is available with `ptdalgorithms` on GitHub.

## 5 Discussion

This paper presents a new open-source library called `ptdalgorithms` written in `C` that implements graph-based algorithms for constructing and reward-transformed continuous and discrete phase-type distributions, and for computing their moments and distribution functions. The computation of moments extends to multivariate phase-type distributions. In addition, the library has a native interface in two programming languages `C` and `R`. Some of the methods presented in this paper build on previously published graph-based matrix manipulation. Still, to our knowledge, this is the first time these graph-based approaches have been applied to phase-type distributions and published as an accessible software package.

The joint distribution function of multivariate discrete phase-type distributions can be described by such a recursive algorithm as we have presented for the marginal distribution function. However, for this application the algorithm must keep track of not just the probability of a vertex being visited but also the rewards accumulated at that point for each vertex. Although computationally infeasible in most situations, such an algorithm would be useful for small state-spaces or accumulated rewards and could be an avenue of possible future work.

Although `ptdalgorithms` provides some support for time-inhomogeneous phase-type distributions, this is achieved by discrete changes to time-homogeneous models. New algorithms are required to provide full support for time-inhomogeneous phase-type distributions in a graph framework.

We have demonstrated that our graph representation is orders of magnitudes faster than the matrix based approach. The straightforward iterative construction of the state space lends itself to powerful modeling, and our algorithms allow the computation of moments and distributions for huge state spaces. The general multivariate phase-type distributions allow marginal expectations and the covariance between events to be studied easily. As `ptdalgorithms` include functions for converting between graph and matrix representations, our library may serve as a plug-in in a multifaceted modeling and inference process. We hope this package will enable users to quickly and accessibly model many complex phenomena in natural sciences, including population genetics.

## Acknowledgements

We are grateful to two anonymous reviewers. Their useful comments and constructive suggestions helped improve a previous version of the manuscript.

